# Allee dynamics: growth, extinction and range expansion

**DOI:** 10.1101/098418

**Authors:** I Bose, M Pal, C Karmakar

## Abstract

In population biology, the Allee dynamics refer to negative growth rates below a critical population density. In this Letter, we study a reaction-diffusion (RD) model of population growth and dispersion in one dimension, which incorporates the Allee effect in both the growth and mortality rates. In the absence of diffusion, the bifurcation diagram displays regions of both finite population density and zero population density, i.e., extinction. The early signatures of the transition to extinction at a bifurcation point are computed in the presence of additive noise. For the full RD model, the existence of travelling wave solutions of the population density is demonstrated. The parameter regimes in which the travelling wave advances (range expansion) and retreats are identified. In the weak Allee regime, the transition from the pushed to the pulled wave is shown as a function of the mortality rate constant. The results obtained are in agreement with the recent experimental observations on budding yeast populations.

---

Biological systems are characterised by dynamics with both local and non-local components. In the case of spatially homogeneous systems, one needs to consider only local dynamics based on reaction/growth kinetics. The concentrations of biomolecules like messenger RNAs (mRNAs) and proteins increase through synthesis and decrease through degradation. The density of a cell population is subjected to changes brought about by birth and death processes. In the latter case, depending upon specific conditions, the cell population acquires a finite density in the course of time or undergoes extinction. In the case of a spatially heterogeneous system, reaction-diffusion (RD) processes govern the dynamics of the system. The RD models have been extensively studied in the context of spatiotemporal pattern formation in a variety of systems [1,2]. One possible consequence of RD processes is the generation of travelling waves which are characteristic of a large number of chemical and biological phenomena [3,4]. The shape of a travelling wave is invariant as a function of time and the speed of propagation is a constant. Biological systems exhibit travelling waves of measurable quantities like biochemical concentration, mechanical deformation, electrical signal, population density etc. One advantage of travelling fronts in biological systems is that for communication over macroscopic distances, the propagation time is much shorter than the time required in the case of purely diffusional processes. In this Letter, we study cell population dynamics in a one-dimensional (1d) spatially heterogeneous system described by the RD equation with the general form

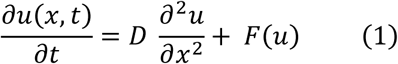

where *u*(*x, t*) denotes the population density at the spatial location *x* and time *t*, D is the diffusion coefficient and the term *F*(*u*) represents the local growth rate incorporating the Allee effect. We derive a number of results on population growth, extinction and range expansion which are in agreement with recent experimental observations in microbial systems.

## 1. Models of Growth and Extinction

In the absence of diffusion, only the local dynamics of the population are relevant and a major issue of interest is the survival or extinction of the population in the steady state. In the case of logistic growth [3],

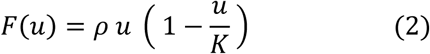

where *ρ* denotes the exponential growth rate and *Κ* the carrying capacity (the population growth stops when *u* = *Κ*). The growth rate *F*(*u*) satisfies the properties

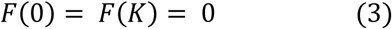

with the steady state *u* = 0 being unstable and the steady state *u* = *Κ* being stable, a case of monostability.

Also,

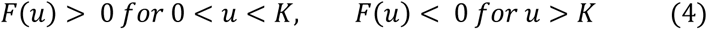

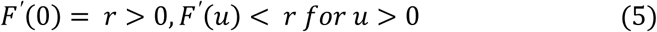

where the prime represents the derivative with respect to *u*. The per capita growth rate, 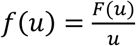, is a monotonically decreasing function of the population density. One common departure from the logistic growth, observed in natural and laboratory populations, is designated as the Allee effect [4,5,6]. In the case of the strong Allee effect, the local growth rate *F*(*u*) is negative when the population density *u* is less than a critical density *β* known as the Allee threshold. The negative growth rate results in population extinction in the long time limit. The most well-studied functional form of *F*(*u*), which illustrates the Allee effect, is

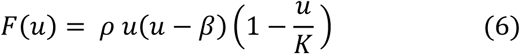

One now has

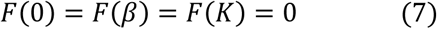

with the steady states *u* = 0 and *u* = *Κ* being stable, a case of bistability, and the state *u* = *β* being unstable. The other conditions on *F*(*u*) are

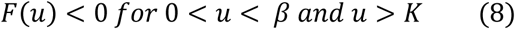

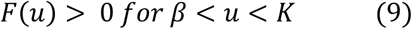

The per capita growth rate *f*(*u*) is negative below the Allee threshold and reaches a maximum value at an intermediate density. In the case of the weak Allee effect, the per capita growth rate is always positive signifying the absence of an Allee threshold. The maximal value again occurs at an intermediate density. A possible origin of the Allee effect lies in the reduced cooperativity amongst the individuals of the population at low population densities [5,6,7]. Examples include mate shortages in sexually reproducing species, less efficient feeding and reduced effectiveness of vigilance and antipredator defences [5,6]. Allee effects have been demonstrated in all major taxonomic groups of animals, in plants [6] as well as in microbial populations[8,9].

There are several ways of modeling the Allee effects [4,10] with most of the models explicitly including the Allee threshold. A less phenomenological model has been proposed by Berec et al. [11] which incorporates the Allee effect in both the birth (reproduction) and death (mortality) processes. The function *F*(*u*) in this case is given by

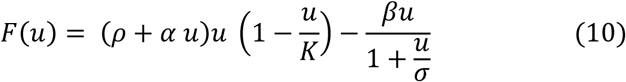

where *ρ* is the basic birth rate, *α* is the Allee effect coefficient with higher values of *α* signifying steeper increases in the reproduction rate, *β* denotes the mortality rate constant when the Allee effect is present, i. e., mortality increases when the population density *u* approaches zero and the parameter *σ* is the population density for which the mortality rate is halved. If *α* = 0 (*σ* = 0), there is no Allee effect on reproduction (mortality). If both the parameters *α* and *σ* are zero, the model reduces to the usual logistic growth model. In Ref. [11], the model studied describes the fragmentation of a population in several patches with the growth kinetics of a subpopulations described by *F*(*u*) in equation (10). The movement of the population between patches occurs through diffusion. The number of patches considered in the study was two and the parameter *α* was taken to be zero. The focus of the study was to investigate the interplay between the Allee effects and collective inter-patch movement. In this Letter, we study the RD model described in equation (1) with the functional form of *F(u)* as given in equation (10). Though the model is of a general nature, our focus in this Letter is on microbial cell populations, motivated by some recent experiments on laboratory populations of budding yeast [8,12,13,14].

## 2. Allee Dynamics

We first report on the steady state properties of the model in the absence of diffusion. Figures 1(a) and 1(b) exhibit the steady state population density *u* versus the mortality rate constant *β* for the cases *α* = 0 and *α* = 1 respectively. In the region of bistability, the two stable steady state branches, represented by solid lines, are separated by a branch of unstable steady states, denoted by a dashed line. The upper stable steady state represents a population with finite density whereas the lower stable steady state (*u* = 0) corresponds to population extinction. The transitions between the bistable and monostable regions occur via the saddle-node (fold) bifurcation at two bifurcation points. In the region of bistability, if the initial population density is above the dotted line, the population acquires a finite density in the steady state. The population undergoes extinction if the initial population density falls below the dotted line. When *α* = 0, the steady states are given by

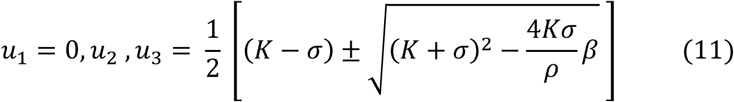

with *u*_1_ and *u*_2_ being stable and *u*_3_ unstable. The region of bistability is obtained in the parameter regime

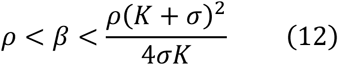

**Figure 1.**
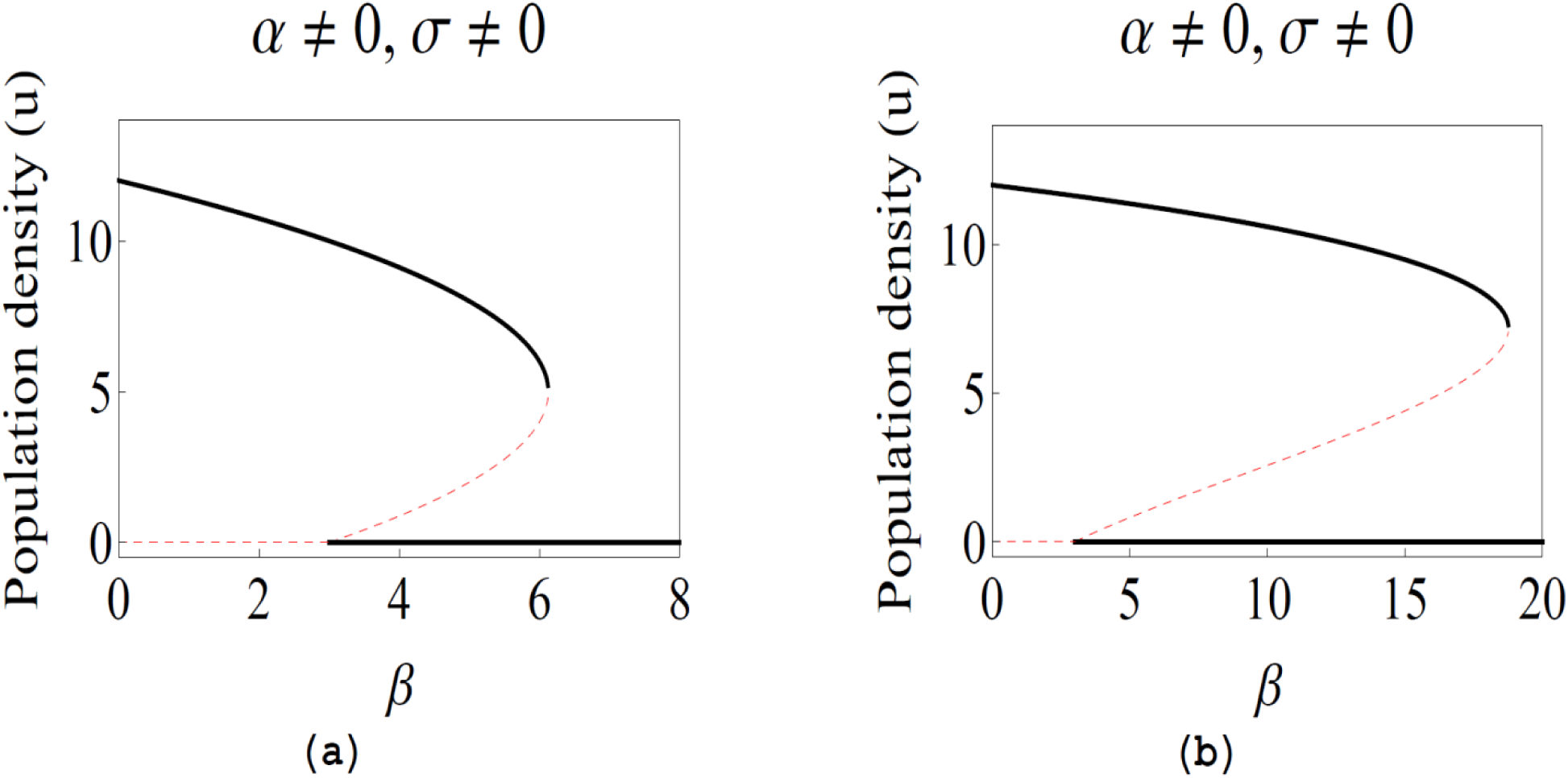
Steady state population density *u* versus the mortality rate constant *β* for the cases (a) *α* = 0 and (b) *α* = 1. The other parameter values are *ρ* = 3, *Κ* = 12, *σ* = 2.

The boundary points of the regime represent the two bifurcation points. When *α* ≠ 0 (figure 1(b)), one obtains the steady states through numerical computation. As depicted in figures 1(a) and 1(b), a tipping point transition occurs at the upper bifurcation point from a finite population density to population collapse.

### 2.1 Early Signatures of Regime Shifts

Recently, a large number of studies have been carried out on the early signatures of tipping point transitions involving sudden regime shifts in systems as diverse as ecosystems, financial markets, population biology, complex diseases, gene expression dynamics and cell differentiation [15,16,17,18]. The early signatures include the critical slowing down and its associated effects, namely, rising variance and the lag-1 autocorrelation function as the bifurcation point is approached. Other signatures have also been proposed, e.g., an increase in the skewness of the steady state probability distribution as the system gets closer to the transition point [15,16]. For our model system (*α* ≠ 0, *β* ≠ 0), we compute the variance and the autocorrelation time (measures the time scale of correlation between the fluctuations at different time points), through simulation of the stochastic difference equation

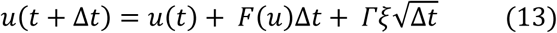

where *F*(*u*) has the form as given in equation (10) and the last term on the right hand side represents the effect of stochasticity (noise). *Γ* is the strength of the additive noise and *ξ* represents a Gaussian random variable with zero mean and unit variance. Let *δu*(*t*) = *u*(*t*) – *u*_2_ represent the deviation of the population density *u* at time *t* from the stable steady state population density *u*_2_. The simulation data are recorded after 1000 time steps (stationarity conditions reached) for an ensemble of five hundred replicate populations. Figure 2A shows the plots of *δu*(*t* + Δ*t*) versus *δu*(*t*) for two distinct values of the bifurcation parameter *β*. The first figure depicts the results far from the bifurcation point with *β* = 6.0 and the second figure corresponds to the value of *β* = 18 which is closer to the bifurcation point *β* = 18.79. The parameter values are Δ*t* = 0.01 and *Γ* = 0.5 with the other parameter values the same as in figure 1(b). Figures 2B and 2C display the autocorrelation time *τ* and the steady state variance *σ*^2^ as a function of the bifurcation parameter *β*. The autocorrelation time *τ* is computed as 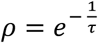 where *ρ* is the lag-1 autocorrelation estimated by the sample Pearson’s correlation coefficient [8].

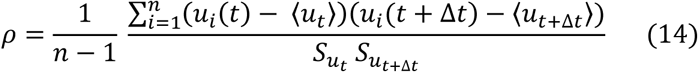

where *i* denotes the sample index, *n* is the total number of samples, 〈*u_t_*〉 is the sample mean and *S_U_t__* the sample standard deviation at time *t*. In the computation, *n* = 500 and the time interval Δ*t* is treated as the unit time interval.

**Figure 2.**
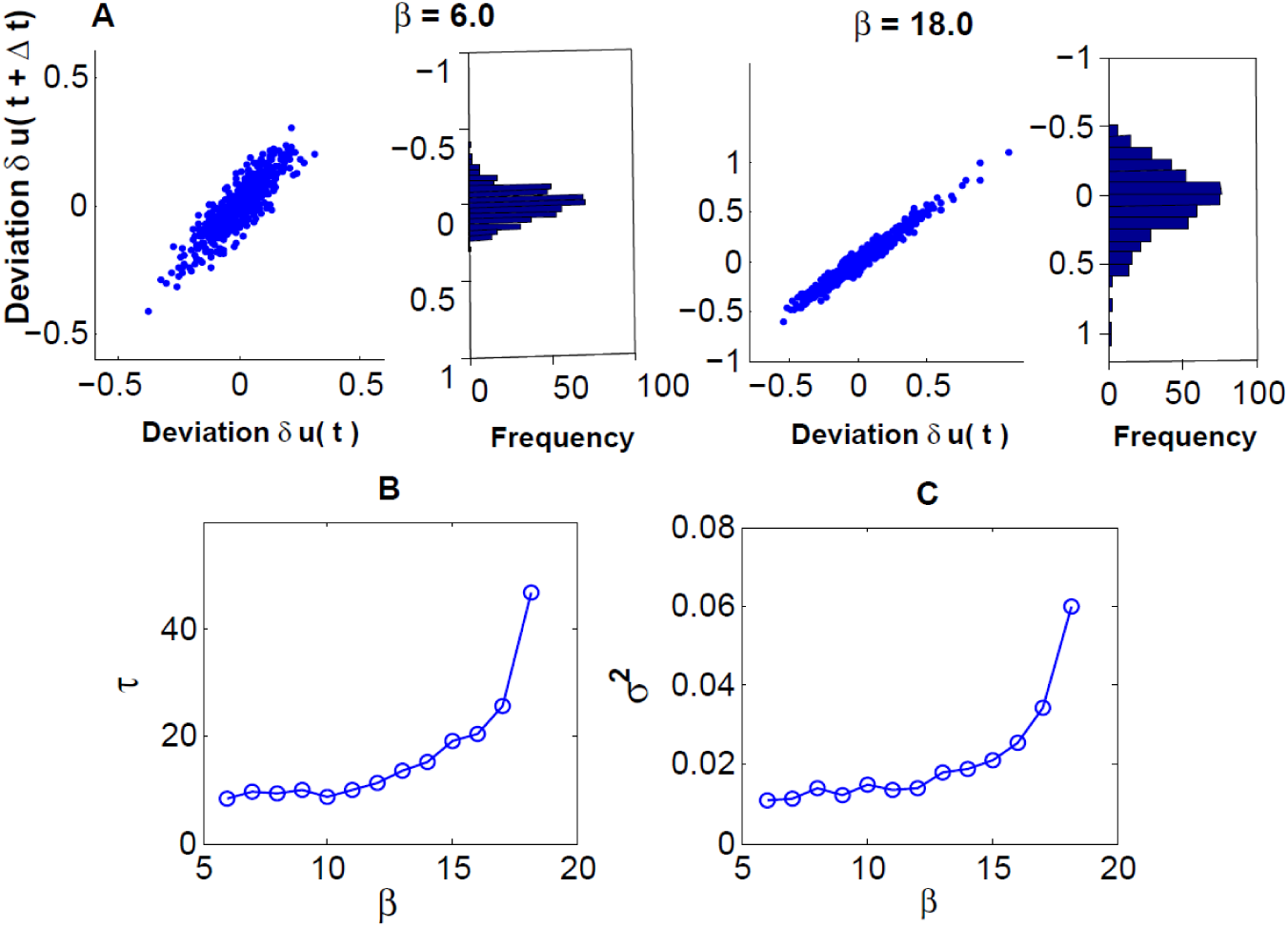
A. Temporal correlations and frequency distributions of deviations from the stable steady state at *β* = 6.0 and *β* = 18.0. B and C. Plots of autocorrelation time *τ* and steady-state variance *σ*^2^ as a function of *β*.

Figure 2 provides numerical evidence that both the variance and the autocorrelation time increase as the bifurcation point is approached. The return time equals the autocorrelation time [8] and its increase in the vicinity of the bifurcation point implies the critical slowing down. The results obtained in our model study are in accordance with the experimental observations of Dai et al. [8] on laboratory populations of budding yeast *S.cerevisiae*. In the experiment, the Allee effect was realized through the cooperative growth of budding yeast in sucrose. Yeast cells are known to hydrolyze sucrose outside the cell thus creating a common pool of hydrolysis products which is utilized by individual cells. The cooperative breakdown of sucrose results in the per capita growth rate becoming maximal at intermediate cell densities, a feature of the Allee effect. Replicate yeast cultures with a wide range of initial cell densities were subjected to daily dilutions into fresh sucrose media. The dilution is equivalent to introducing mortality in the population with higher dilution factors implying larger mortality rates. An experimental bifurcation diagram depicting the population density versus the dilution factor, which serves as the bifurcation parameter, was mapped out. The bifurcation was of the saddle-node type and a tipping point transition from a population of finite density to population extinction was identified. A simple two-phase growth model provided a good fit to the experimental data. In the model, the population growth was slow exponential at low densities and faster logistic at larger densities, based on experimental evidence. The theoretically predicted early warning signals of the tipping point transition were also tested experimentally. The experimentally obtained bifurcation diagram and the early warning signatures (figures 1(E) and figure 3 of Ref. [8]) are qualitatively similar to the theoretical plots (figures 1 and 2) computed in our model study. The model serves as a microscopic model to describe population growth and extinction in the budding yeast population and explicitly includes the parameters describing growth, mortality and the Allee effect.

## 3. Reaction-Diffusion Model with Allee Dynamics

We next consider the full RD model shown in equation (1). Depending on the form of *F*(*u*), two prototype cases are possible, monostability and bistability. The RD equation, with *F*(*u*) given in equation (2) (logistic growth), was first considered by Fisher [19] to describe the spread of a favourable gene in a population. The equation describes monostability and is referred to as the Fisher-Kolmogorov, Petrovskii and Piskunov (FKPP) equation. When *F*(*u*) is of the form given in equation (6) and the conditions in equations (8) and (9) hold true, i.e., the strong Allee effect is considered, the situation is that of bistability. It has been rigorously shown [4,20] that for suitable initial conditions *u*(*x*, 0), the disturbance evolves into a monotonic travelling front *u* = *q*(*x – ct*), with constant speed *c*, joining the stable steady state with finite population density to the steady state *u* = 0 (unstable in the case of monostability and stable in the case of bistability). Populations occupy new territory through a combination of local growth and diffusion. If the population front moves from a region of finite *u* to one with *u* = 0, the phenomenon is characterised as range expansion or invasion. If the travelling front moves in the opposite direction, the population retreats rather than advances.

In the case of the FKPP equation describing generalized logistic growth, a travelling wave solution with speed *c* satisfies the relation *c* ≥ *c_min_* with

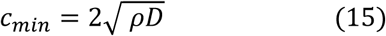

It has been shown that in the case of a compact initial condition, the initial population distribution always develops into a travelling population front with the minimum speed *c* = *c_min_*. The scaling 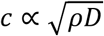 is known as the Luther formula [21] and holds true over many orders of magnitude in many chemical and biological systems [22]. A recent example pertains to the propagation of gene expression fronts in a one-dimensional coupled system of artificial cells [22]. We next consider the case of reaction-diffusion incorporating the Allee effect. When strong Allee effect conditions prevail, i.e., 0 < *β* < *Κ* in equation (6), the travelling wave front has a unique speed given by [3,10, 20]

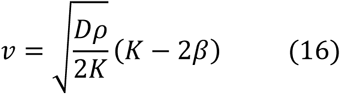

In contrast to the case of logistic growth in which the travelling front has a finite value for the minimum speed, the velocity 𝑣 becomes zero at the so-called Maxwell point 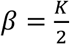. Also, the front has a unique speed instead of a spectrum of possible values above or equal to a minimum speed. If *β* and *Κ* are treated as general parameters without any specific physical interpretation, then the parameter regions 0 < *β* < *Κ* and – *Κ* < *β* < 0 are associated with the strong and weak Allee effects respectively. For *β* ≤ –*Κ*, the Allee effect is absent and the growth rate is of the generalized logistic type.

In the case of the FKPP equation, the minimal speed of the travelling front can be written as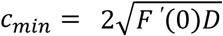. The travelling wave is designated as the pulled wave since the wave is pulled by the leading edge of the population distribution [14]. The solution with the minimal speed defines the critical front while the solutions for which the wave speed c is > *c_min_* define the super-critical fronts. A pulled wave is either a critical front or any super-critical front. In the case of a pushed wave, the minimal speed satisfies the relation

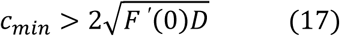

The speed of the travelling front is now determined by the whole front rather than by only the leading edge. In the monostable case ( logistic growth and weak Allee effect), the travelling front can be classified as either pulled or pushed wave whereas in the case of bistability (strong Allee effect), the travelling wave is always a pushed wave with a unique speed greater than 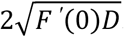. The theoretical prediction of a pulled to pushed wave transition [14] has recently been verified by Gandhi et al. in a laboratory population of budding yeast as the cooperativity in growth was increased. The transition occurred at some intermediate magnitude of the Allee effect in the regime corresponding to the weak Allee effect.

The RD model studied in this letter with *F*(*u*) as given in equation (10), has a more general form than the model, most often studied, in the context of the Allee effect (*F*(*u*) as given in equation (6)). The existence of a travelling wave solution in this model is supported by the theorem proposed by Fife and McLeod [23]. The theorem states that the RD equation (1), with (*F*(*u*_1_)*F*(*u*_2_)*F*(*u*_3_) = 0 and *F′*(*u*_1_) < 0, *F′* (*u*_2_) < 0, *F′* (*u*_3_) > 0, has a travelling wave solution *u*(*x,t*) = *U*(*x* – 𝑣*t*) for exactly one value of the wave speed 𝑣. The solutions *u*_1_ and *u*_2_ describe stable steady states and the solution *u*_3_ an unstable steady state. Also, *U*(−∞) = *u*_2_ and *U*(+∞) = *u*_1_, i.e., the travelling wave connects the solutions *u*_1_ and *u*_2_. We assume that the parameter *α* = 0 in equation (10), i.e., the Allee effect is present only in the mortality rate. In this case, the analytic solutions for the steady states are known (equation (11)) and one can look for travelling wave solutions for the RD equation. In the following, we report on the results obtained from numerical computations.

### 3.1 Range Expansion, Pushed and Pulled waves

The 1d RD system has boundaries located at −*L* and +*L* respectively. We solve the RD equation on Mathematica with the compact initial conditions

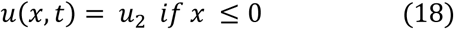

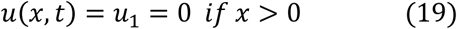

The value of *L* is chosen to be 50, the diffusion coefficient *D* = 1 and the other parameter values are the same as in figure 1(a). Figure 3 shows the computed travelling wave solutions at the time points *t* = 20,40,60 and 80 and *β* = 5.

**Figure 3.**
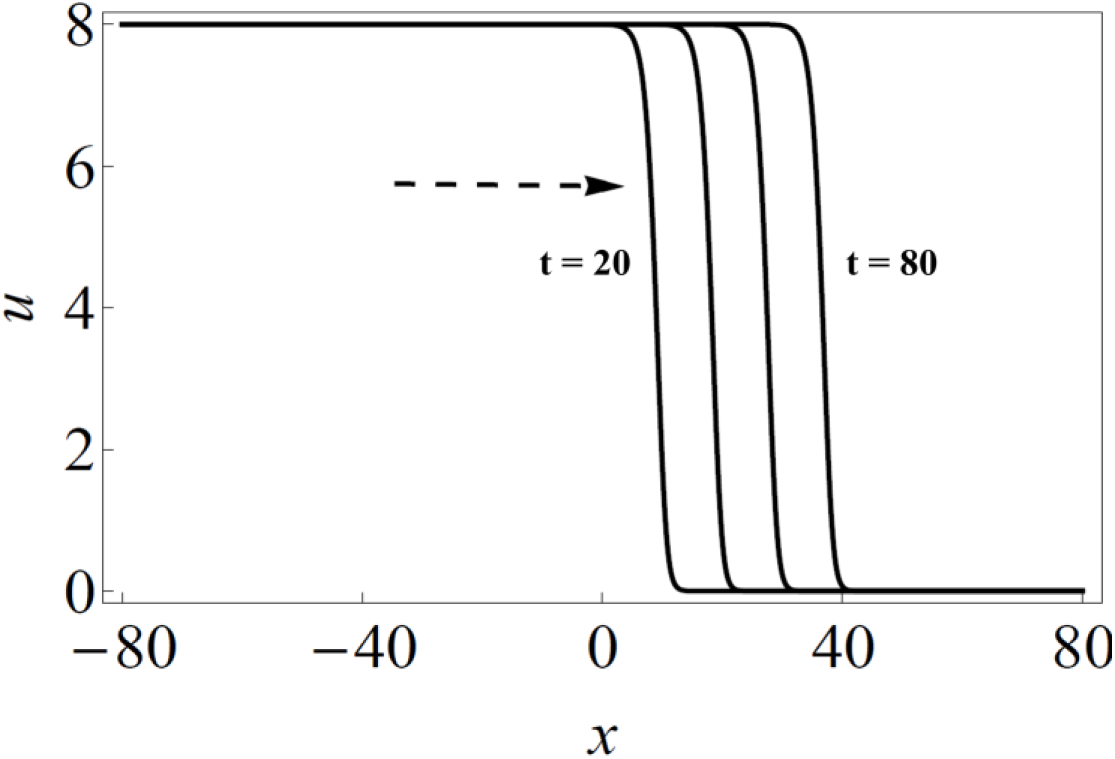
Travelling wave solutions of the RD equation (1) with *F*(*u*) given in equation (10) for *α* = 0. The solutions are computed at the time points *t* = 20, 40, 60 and 80 The other parameter values are *D* = 1, *ρ* = 3, *Κ* = 12, *σ* = 2 and *β* = 5.

The front moves from the left to the right with the population of finite density, *u*_2_, advancing into the empty region, *u*_1_ = 0. Figure 4 shows the speed 𝑣 of the travelling wave versus the mortality rate parameter *β* ( solid circles). The speed was computed by plotting the midpoint *x_m_* of the density profile versus time *t* and determining the slope of the straight-line fit [13]. The open circles represent the speed given by the formula, 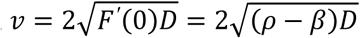 which corresponds to the speed of a pushed wave. The computed speed of the travelling wave and that of the pulled wave are nearly equal up to a crossover point denoted by a big circle. The RD equation reduces to the FKPP equation when the parameter *β* = 0. In the case of compact initial conditions, as given in equations (18) and (19), the exact expression for the travelling wave velocity is that of the pushed wave. The small difference with the computed value can be ascribed to the finite size of the system and the small but finite errors associated with the numerical computation of speed. For *β* < *ρ* = 3, the system is monostable corresponding to the weak Allee effect regime. The strong Allee effect regime is bistable with the parameter range as specified in equation (12). At the crossover point, *β* ≈ 1.823, the travelling wave changes its nature from pulled to pushed wave. The speed of the travelling wave becomes zero at the Maxwell point *β_MP_* ≈ 5.6259. The travelling wave then reverses its direction, i.e., the wave retreats from the empty region rather than advance into it. The situation continues till the upper bifurcation point, 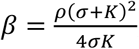 marking the region of bistability. The transition from the pulled to the pushed wave occurs in the weak Allee effect regime, in agreement with the experimental observation of the transition in a laboratory population of budding yeast [14].

**Figure 4.**
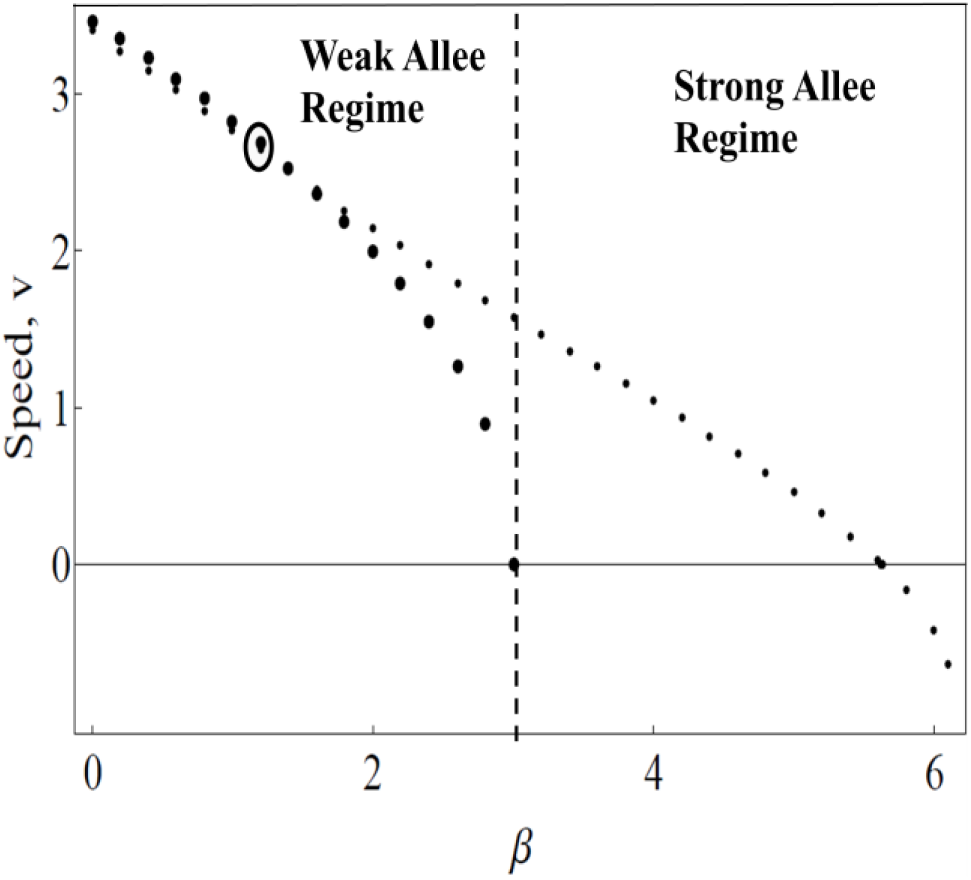
Speed 𝑣 versus *β*. The big open circle denotes the cross-over region from pushed to pulled wave.

## 4. Conclusion

In this Letter, we have studied a RD model in 1d with the Allee effect incorporated in the dynamics of the model. The model in its full form has not been studied previously. The strength of the model lies in the fact that the growth and mortality rate constants appear explicitly, as parameters, in the model which are experimentally controllable. The model is general in nature and applicable for the study of population dynamics based on growth and diffusion. Though we could not derive analytical results for the full model, the results obtained through numerical computations are in agreement with the experimental observations on budding yeast populations. Recent advances in cancer research have drawn parallels between tumour and population dynamics [7]. There is some experimental evidence [7] that the Allee effect forms an important component of the tumour dynamics. The RD model studied by us is expected to be relevant in capturing some essential features of tumour growth and spread.

## Acknowledgements

IB acknowledges the support by CSIR, India, vide sanction Lett. No. 21 (0956)/13-EMR-II dated 28.04.2014. CK acknowledges the support by National Network for Mathematical and Computational Biology, India for carrying out part of the study. MP acknowledges support from Bose Institute, India for carrying out the research study.

